# BRET-based self-cleaving biosensors for SARS-CoV-2 3CLpro Inhibitor Discovery

**DOI:** 10.1101/2021.07.28.454072

**Authors:** Ningke Hou, Chen Peng, Lijing Zhang, Yuyao Zhu, Qi Hu

## Abstract

The 3C-like protease (3CLpro) of SARS-CoV-2 is an attractive drug target for developing antivirals against SARS-CoV-2. A few small molecule inhibitors of 3CLpro are in clinical trials for COVID-19 treatments and more inhibitors are being developed. One limiting factor for 3CLpro inhibitors development is that the cellular activities of such inhibitors have to be evaluated in a Biosafety Level 3 (BSL-3) or BSL-4 laboratory. Here, we design genetically encoded biosensors that can be used in BSL-2 laboratories to set up cell-based assays for 3CLpro inhibitor discovery. The biosensors were constructed by linking a green fluorescent protein (GFP2) to the N-terminus and a Renilla luciferase (RLuc8) to the C-terminus of SARS-CoV-2 3CLpro, with the linkers derived from the cleavage sequences of 3CLpro. After over-expression of the biosensors in HEK293 cells, 3CLpro can be released from GFP2 and RLuc by self-cleavage, resulting in a decrease of the bioluminescence resonance energy transfer (BRET) signal. Using one of these biosensors, pBRET-10, we evaluated the cellular activities of several 3CLpro inhibitors. These inhibitors restored the BRET signal by blocking the proteolysis of pBRET-10, and their relative activities measured using pBRET-10 were consistent with their anti-SARS-CoV-2 activities reported previously. We conclude that the biosensor pBRET-10 is a useful tool for SARS-CoV-2 3CLpro inhibitor discovery. Furthermore, our strategy can be used to design biosensors for other viral proteases that share the same activation mechanism as 3CLpro, such as HIV protease PR and HCV protease NS3.

**Highlights:**

- Sensitive cell-based biosensors for 3CLpro inhibitor discovery in BSL-2 laboratories.
- The BRET-based self-cleaving biosensors mimic the *in vivo* autoproteolytic activation of 3CLpro.
- Similar biosensors can be designed for other self-cleaving proteases, such as HIV protease PR and HCV protease NS3.

## 1. Introduction

The severe acute respiratory syndrome coronavirus 2 (SARS-CoV-2) causing the global pandemic of COVID-19 poses a great threat to public health (Zhou et al., 2020). Despite several vaccines have been accessible, effective antivirals are still urgently needed for the treatment of COVID-19 (Grobler et al., 2020). The RNA genome of SARS-CoV-2 encodes two large overlapping polyproteins pp1a and pp1ab, and several structural proteins and accessory proteins (Hartenian et al., 2020). During virus replication in host cells, pp1a and pp1ab are expressed and then cleaved to generate 16 non-structural proteins (nsps). The cleavages are catalyzed by nsp3 and nsp5 – two proteases included in the 16 nsps. Specifically, the papain-like protease (PLpro) domain of nsp3 cleaves the peptides bonds between nsp1 and 2, nsp2 and 3, and nsp3 and 4; the peptide bonds between other nsps are cleaved by nsp5 (also called 3C-like protease, 3CLpro or the main protease). Inhibition of 3CLpro is an effective strategy to develop antivirals against SARS-CoV-2 (Zumla et al., 2016). Several 3CLpro inhibitors have been reported, two of which are now in COVID-19 clinical trials (Dai et al., 2020; de Vries et al., 2021; Drayman et al., 2021; Fu et al., 2020; Jin et al., 2020; Vandyck and Deval, 2021; Zhang et al., 2020).

Enzymatic assays using purified 3CLpro were frequently used in the initial screening of 3CLpro inhibitors, but to evaluate the cell permeability and cellular activities of the inhibitors, cell-based antiviral assays are necessary. The requirement of Biosafety Level 3 (BSL-3) or BSL-4 laboratories for doing the cell-based anti-SARS-CoV-2 assays has slowed the development of 3CLpro inhibitors.

To set up cell-based 3CLpro assays that can be done in BSL-2 laboratories, several biosensors have been developed. The first is called Flip-GFP that has a 3CLpro cleavage site inserted into GFP; the fluorescence of GFP was decreased by the insertion but could be restored by the 3CLpro-catalyzed cleavage (Froggatt et al., 2020). Similar biosensors have been developed by other groups (O’Brien et al., 2021; Gerber et al., 2021). The second is an engineered luciferase having two complementary luciferase fragments linked by a 3CLpro cleavage site; the luminescence was lost by 3CLpro cleavage and restored when the 3CLpro activity was inhibited (Rawson et al., 2021). The third is a GFP fusion protein having an ER targeting domain linked to the C-terminus of GFP through a 3CLpro cleavage site; 3CLpro-catalyzed cleavage led to a translocation of the GFP from ER to the nucleus which can be quantified using light microscopy (Pahmeier et al., 2021). The fourth is a biosensor in which a Src myristoylation domain and a HIV-1 Tat-GFP fusion protein are linked to the N- and C-terminus of 3CLpro, respectively, through 3CLpro cleavage sites; expression of this biosensor in HEK 293T cells showed little GFP fluorescence while inhibition of 3CLpro greatly increased the GFP fluorescence, probably because 3CLpro-catalyzed self-cleavage led to degradation of the biosensor (Moghadasi et al., 2020).

The limitations of these biosensors are that (1) their readouts are highly dependent on the expression level of the biosensors, and for the first three types of biosensors, the expression level of 3CLpro also affects the readouts; (2) the first three types of biosensors require either co-transfection of two plasmids (the biosensor and 3CLpro plasmids) or transfection of 3CLpro into cells stably expressing the biosensors, while the fourth only needs to transfect one plasmid but its sensitivity to 3CLpro inhibitor GC376 was much lower.

In this study, we developed a series of BRET-based biosensors to set up cell-based assays for 3CLpro inhibitor discovery. We linked a green fluorescent protein (GFP2) and a Renilla luciferase (RLuc8) to the N- and C-terminus of SARS-CoV-2 3CLpro, respectively, using 3CLpro cleavage sequences as the linkers (Fig. 1A&B). The BRET from Rluc8 to GFP2 of the biosensors was disrupted upon self-cleavage catalyzed by 3CLpro and can be restored by adding 3CLpro inhibitors. The effect of variance in the biosensor expression level on the readouts was minimized by normalizing the BRET signal with the luminescent signal of RLuc8.

**Figure 1.**
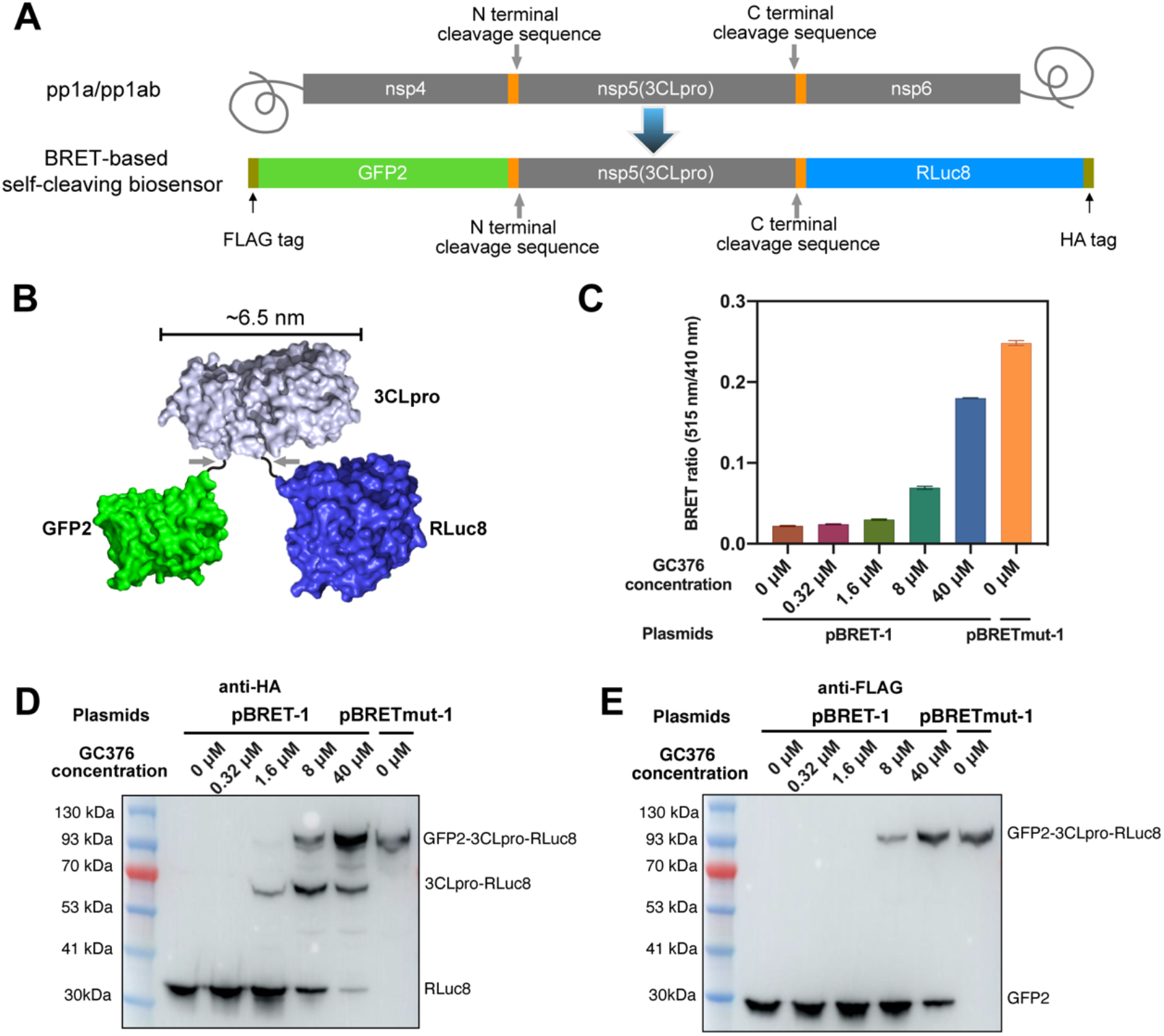
A) The domain organization of SARS-CoV-2 polyproteins (pp1a or pp1ab) (top) and the BRET-based self-cleaving biosensor (bottom). B) An illustration of the BRET-based self-cleaving biosensor showing the relative position of GFP2, 3Clpro and RLuc8. C) The BRET ratio (GFP_515 nm_/Rluc_410 nm_) of HEK 293T cells 24 hours post-transfection of the plasmid carrying biosensor pBRET-1. The 3CLpro inhibitor GC376 at the indicated working concentrations were added right after transfection. The biosensor with a C145A mutation in 3CLpro (pBRET_mut_-1) was used as non-cleavable control. The data represent the mean ± SD of three independent measurements. D and E) The self-cleavage of pBRET-1 in the presence of indicated concentrations of GC376 was detected by Western blot analysis using an anti-HA antibody (D) or an anti-FLAG antibody (E).

## 2. Material and methods

### 2.1. Construction of plasmids

The gene sequences of RLuc8 and GFP2 are the same as that reported previously (Bery et al., 2018). The gene sequences of SARS-CoV-2 3CLpro and its cleavage sites are the same as that in the SARS-CoV-2 genome (NC_045512.2). The DNA fragment encoding the biosensor pBRET-1 was synthesized at GENEWIZ (Suzhou, China) and inserted into pcDNA3.1 vector at the site after the FLAG-tag. Then an HA-tag was added to the C-terminus of pBRET-1 by Gibson homologous recombination using primers HA-F and HA-R (Table S1). The plasmid of pBRET_mut_-1 was constructed by introducing the 3CLpro C145A mutation into pBRET-1 through site-directed mutagenesis using primers C145A-F and C145A-R (Table S1). The plasmids carrying other pBRET biosensors were constructed on the basis of pBRET-1 using Gibson homologous recombination method. The primers were shown in Table S1. The protein sequences of all the BRET-based self-cleaving biosensors were shown in Table S2.

### 2.2. Cell culture

HEK 293T cells were cultured in a humidified incubator maintained at 37 °C with 5% CO_2_, using the high-glucose Dulbecco’s modified Eagle’s medium (Gibco) supplemented with 100 U/mL penicillium-streptomycin (HyClone) and 10% fetal bovine serum (Gibco).

### 2.3. 3CLpro inhibitors

GC376 (Selleck, S0475) and Boceprevir (Selleck, S3733) were purchased form Selleck. Compounds 11a and 13b were synthesized following protocols reported previously (Dai et al., 2020; Zhang et al., 2020).

### 2.4. Western blot and BRET assays

To monitor the self-cleavage of pBRET-1 and to evaluate its sensitivity to 3CLpro inhibitor GC376, HEK 293T cells were seeded into 6-well cell culture plates at about 40% confluence, and after 24 hours the cells were transfected with plasmids carrying pBRET-1 (5 μg/well) using PEI as the transfection reagent. After transfection, 3CLpro inhibitors in DMSO were added into the cell culture to reach the indicated working concentrations. The final DMSO concentration in the cell culture was 0.5%. After additional 24 hours, the cells were harvested and equally divided into two parts: one part for western blot analysis, and the other for BRET assay.

For western blot analysis, HEK 293T cells were lysed using RIPA buffer (Beyotime, P0013B). Then equal amounts of total protein in each condition were resolved by SDS-PAGE and transferred to a PVDF membrane (Merck Millipore, ISEQ00010). The PVDF membrane was blocked with 1% (for anti-FLAG antibody) or 3% (for anti-HA antibody) non-fat milk in TBST buffer (Tris-buffered saline containing 0.05% Tween-20) overnight at 4 °C and incubated with anti-FLAG antibody (Sigma, F1804, 1:1000 dilution) or anti-HA antibody (Abcam, ab9110, 1:1000 dilution) for 2 hours at room temperature. After being washed three times with TBST buffer, the PVDF membrane was incubated with anti-rabbit IgG (Merck Millipore, AP156P, 1:20000 dilution) or anti-mouse IgG (Merck Millipore, AP127P; 1:20000 dilution) for 1 hours, washed with TBST buffer three times, and developed using Pierce ECL western blotting substrate (CWBIO, CW0049S).

For BRET assay, the cells were resuspended in ice-cold PBS and transferred to a 96-well clear-bottom white plate (Corning, 3610). After adding coelenterazine 400a (Cayman, 16157) to a final concentration of 20 μM, the luminous signal at 410 nm and fluorescent signal at 515 nm were measured using a BioTek microplate reader (Biotek Synergy NEO2). The BRET ratio was calculated using the following equation:

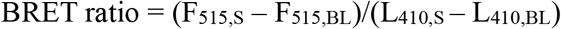

in which F_515,S_ and L_410,S_ are the fluorescent (515 nm) and luminescent signals (410 nm), respectively, of cells expressing pBRET biosensors, and F_515,BL_ and L_410,BL_ are the fluorescent (515 nm) and luminescent signals (410 nm), respectively, of HEK 293T cells without pBRET biosensors.

To set up high-throughput BRET assay for optimizing pBRET-1 and evaluating the activities of different 3CLpro inhibitors, HEK 293T cells were seed into a 96-well clear-bottom white plate (Corning, 3610) at about 40% confluence, and after 24 hours the cells were transfected with plasmids carrying the biosensors (0.4 μg/well) using PEI as the transfection reagent. Then 3CLpro inhibitors in DMSO were added into the cell culture to reach the indicated working concentrations. Twenty-four hours later, the BRET ratios were measured using same protocols as described above.

## 3. Results and discussion

### 3.1. Design of a BRET-based self-cleaving biosensor, pBRET-1

The maximal distance for BRET is about 10 nm (Bacart et al., 2008). According to a crystal structure of SARS-CoV-2 3CLpro (PDB code: 6Y2E), the distance between the N- and C-terminus of 3CLpro is 22.9 Å (Zhang et al., 2020). To construct our first biosensor — pBRET-1, we linked GFP2 to the N-terminus of SARS-CoV-2 3CLpro using the cleavage sequence between nsp4 and 3CLpro as the linker, and linked RLuc8 to the C-terminus of 3CLpro using the cleavage sequence between 3CLpro and nsp6 as the linker; furthermore, we added a FLAG tag before GFP2 and a HA tag after RLuc8 (Fig. 1A). We also constructed pBRET_mut_-1, in which the catalytic residue C145 of 3CLpro was mutated to alanine.

Transient expression of pBRET-1 in HEK293 cells resulted in a BRET ratio (see the methods) of about 0.02, while transient expression of pBRET_mut_-1 showed a BRET ratio of 0.25 (Fig. 1C). Adding GC376, a reported inhibitor of 3CLpro (Fu et al., 2020), to the cell culture increased the BRET ratio of pBRET-1 in a concentration-dependent manner. These results indicate that the BRET ratio of pBRET-1 is negatively associated with the protease activity of 3CLpro.

We also monitored the self-cleavage of pBRET-1 in HEK 293T cells using Western blot. We first used anti-HA antibody to detect the self-cleavage products (Fig. 1D). For pBRET-1 in the absence of GC376, only RLuc was detected; as the concentration of GC376 increased, the band of the 3CLpro-RLuc8 fragment and that of the full-length GFP2-3CLpro-Rluc8 fusion protein appeared. We also used anti-FLAG antibody to detect the self-cleavage products (Fig. 1E). Interestingly, as the concentration of GC376 increased, only the bands of GFP2 and the full-length GFP2-3CLpro-Rluc8 fusion protein were detected, but no band of the GFP2-3CLpro fragment. These results suggest that the cleavage at the N-terminus of 3CLpro occurred before the cleavage at the C-terminus, which is consistent with the maturation process of 3CLpro reported previously (Li et al., 2010).

### 3.2. Optimization of pBRET-1

There are eleven 3CLpro cleavage sites in the polyproteins pp1a and pp1ab of SARS-CoV-2 (Mody et al., 2021). The efficiencies of 3CLpro to cleave these sites are different. We presume that modulating the self-cleaving efficiency of pBRET-1 by changing the cleavage sequence between GFP2 and 3CLpro and that between 3CLpro and RLuc8 may increase the sensitivity of the biosensor to 3CLpro inhibitors. As shown in Fig. 2, nine pBRET biosensors were designed and their sensitivities to GC376 were tested. Among them, pBRET-10 showed the highest sensitivity, with an EC_50_ value of 2.72 μM for GC376; in contrast, the EC_50_ value measured using pBRET-1 was 11.60 μM (Fig. 2C).

**Figure 2.**
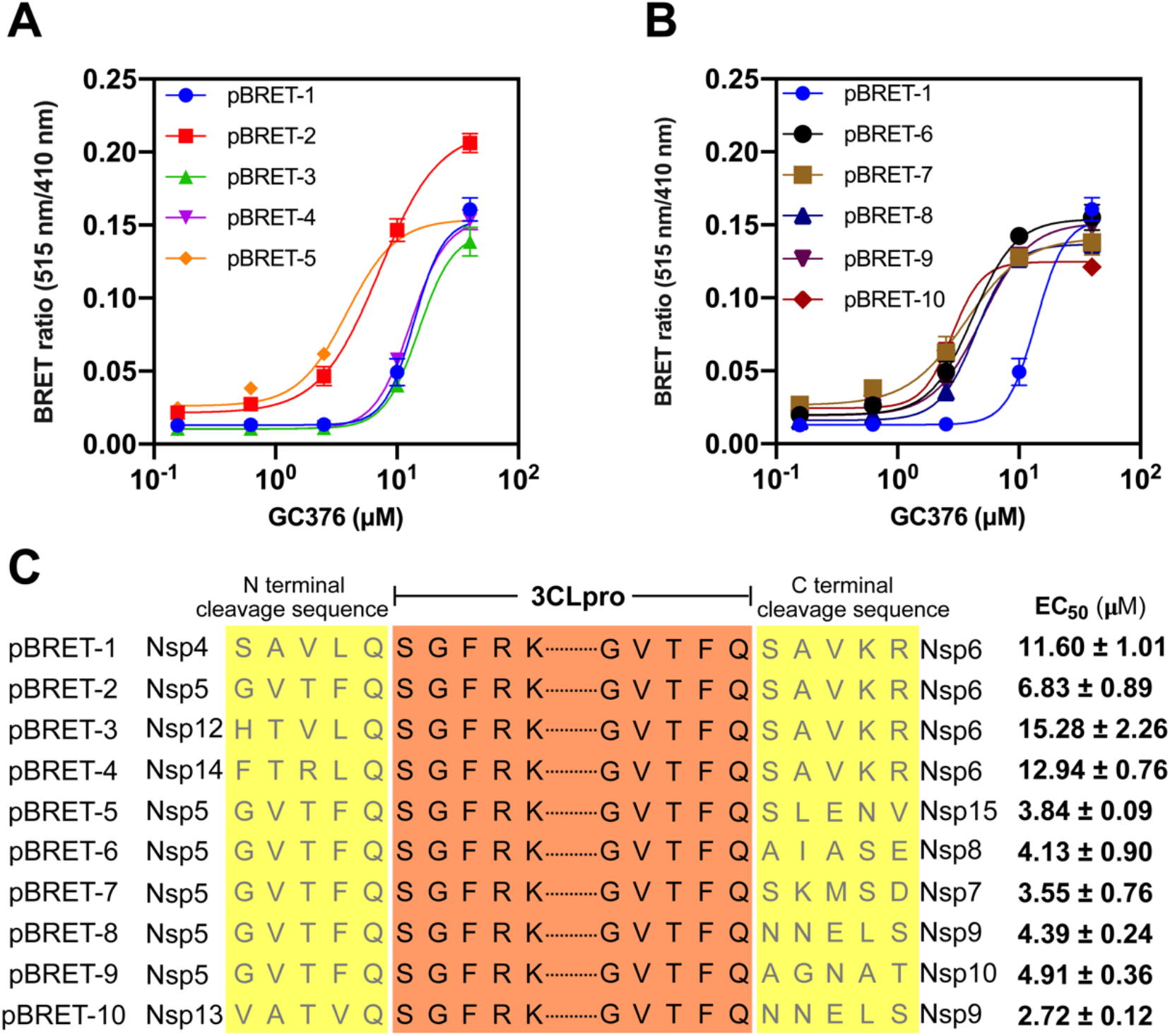
A and B) The BRET ratio (GFP_515 nm_/RLuc_410 nm_) of HEK 293T cells 24 hours post-transfection of different biosensor plasmids (pBRET-1 to pBRET-10). The 3CLpro inhibitor GC376 was diluted into cell culture media at the indicated working concentrations right after transfection. The data represent the mean ± SD of three independent measurements. C) The amino acid sequences of the 3CLpro cleavage sites at the N- and C terminus of the ten biosensors, and the EC_50_ of GC376 measured using these biosensors.

### 3.3. Evaluation of the cellular activities of 3CLpro inhibitors using pBRET-10

As pBRET-10 has the highest sensitivity, we measured the EC_50_ values of three other 3CLpro inhibitors (Boceprevir, compounds 11a and 13b) using pBRET-10 (Table 1) (Dai et al., 2020; Fu et al., 2020; Zhang et al., 2020). Compound 11a showed an activity slightly lower than GC376, while the activities of Boceprevir and compound 13b were an order of magnitude lower than that of GC376. The EC_50_ values of GC376 and Boceprevir are comparable to that measured using cell-based anti-SARS-CoV-2 assays (Fu et al., 2020). But for compounds 11a and 13b, the EC_50_ values from our measurement were about 9 times the reported values from anti-SARS-CoV-2 assays (Dai et al., 2020; Zhang et al., 2020).

**Table 1.** The EC_50_ values of four reported 3CLpro inhibitors measured using pBRET-10 and the corresponding EC_50_ values from anti-SARS-CoV-2 assays.

| 3CLpro inhibitors | EC<sub>50</sub> measured using pBRET-10 (μM) | Reported EC<sub>50</sub> against SARS-CoV-2 (μM) | References |
| --- | --- | --- | --- |
| GC376 | 2.72 ± 0.12 | 0.70 (Vero E6 cells, MOI 0.01);<br>2.20 (Vero E6 cells, MOI 0.01) | (Fu et al., 2020);<br>(Froggatt et al., 2020) |
| Boceprevir | 36.31 ± 2.39 | 15.57 (Vero E6 cells, MOI 0.01) | (Fu et al., 2020) |
| 11a | 4.59 ± 0.23 | 0.53 (Vero E6 cells, MOI 0.05) | (Dai et al., 2020) |
| 13b | 37.11 ± 1.91 | 4~5 (Calu-3 cells, MOI 0.05) | (Zhang et al., 2020) |

## 4. Conclusion

We have developed a class of BRET-based self-cleaving biosensors that can be used in BSL-2 laboratories to set up cell-based assays for 3CLpro inhibitor discovery. One of them, pBRET-10, showed comparable sensitivity to cell-based antiviral assays. Self-cleavage catalyzed by 3CLpro in these biosensors mimics the activation process of 3CLpro during coronavirus replication. In addition to the 3CLpro of SARS-CoV-2, similar biosensors can be developed to screen inhibitors of 3C-like proteases of other coronaviruses. Furthermore, many other viruses (such as HIV, HCV, Dengur virus, Zika virus, and West Nile virus) also utilize a replication strategy involving the expression of a polyprotein containing a self-cleaving protease (Huang et al., 2019; Lin, 2006; Majerová et al., 2019; Suthar et al., 2013; Yost and Marcotrigiano, 2013); therefore, our strategy can also be used to develop biosensors for proteases of these viruses, for example, the HIV protease PR and HCV protease NS3.

## Declaration of competing interest

The authors declare the following financial interests which may be considered as potential competing interests: One or more authors have a pending patent related to this work.

## CRediT authorship contribution statement

**Ningke Hou:** Conceptualization, Methodology, Investigation, Data curation, Visualization, Writing – original draft. **Chen Peng:** Investigation, Data curation, Writing – original draft. **Lijing Zhang:** Resources, Validation. **Yuyao Zhu:** Investigation. **Qi Hu:** Supervision, Conceptualization, Project administration, Funding acquisition, Writing – Review & Editing.

## Acknowledgements

The work is supported by Westlake Education Foundation and Tencent Foundation.

