## Supplementary material for "BRET-based self-cleaving biosensors for SARS-CoV-2 3CLpro Inhibitor Discovery": Table S1 and Table S2

This supplementary material file contains:

Table S1. The primer sequences used in construction of plasmids of BRET-based self-cleaving biosensors.

Table S2. The protein sequences of BRET-based self-cleaving biosensors.

Table S1. The primer sequences used in construction of plasmids of BRET-based self-cleaving biosensors.

| Primer name | Sequences |
| --- | --- |
| HA-F | 5’-ACCCATACGATGTTCCAGATTACGCTTAATCTAGAGGGCCCGTTTAAACCCG-3’ |
| HA-R | 5’-TGGAACATCGTATGGGTAAGATCCCTCGAGCTGCTCGTTCTTCAGCAC-3’ |
| C145A-F | 5’-GGTTCAGCTGGTAGTGTTGGTTTTAACATAGATTATGACTGTGTCTC-3’ |
| C145A-R | 5’-ACACTACCAGCTGAACCATTAAGGAATGAACCCTTAATAGTGAAATTG-3’ |
| pBRET-2-F | 5’-GGTGTTACTTTCCAAAGTGGTTTTAGAAAAATGGCATTCCCATCTGG-3’ |
| pBRRT-2-R | TTGGAAAGTAACACCAGATCCTCCGCCGCCCTTG-3’ |
| pBRET-3-F | CATACAGTCTTACAGAGTGGTTTTAGAAAAATGGCATTCCCATCTGG-3’ |
| pBRET-3-R | CTGTAAGACTGTATGAGATCCTCCGCCGCCCTTG-3’ |
| pBRET-4-F | TTTACAAGACTTCAGAGTGGTTTTAGAAAAATGGCATTCCCATCTGG-3’ |
| pBRET-4-R | CTGAAGTCTTGTAAAAGATCCTCCGCCGCCCTTG-3’ |
| pBRET-5-F | AGTTTAGAAAATGTGGGCGGCGGAGGATCTATG-3’ |
| pBRET-5-R | CACATTTTCTAAACTTTGGAAAGTAACACCTGAGCATTGTC-3’ |
| pBRET-6-F | GCTATAGCCTCAGAGGGCGGCGGAGGATCTATGAC-3’ |
| pBRET-6-R | CTCTGAGGCTATAGCTTGGAAAGTAACACCTGAGCATTGTCTAAC-3’ |
| pBRET-7-F | TCTAAAATGTCAGATGGCGGCGGAGGATCTATGAC-3’ |
| pBRET-7-R | ATCTGACATTTTAGATTGGAAAGTAACACCTGAGCATTGTCTAAC-3’ |
| pBRET-8-F | AATAATGAGCTTAGTGGCGGCGGAGGATCTATGAC-3’ |
| pBRET-8-R | ACTAAGCTCATTATTTTGGAAAGTAACACCTGAGCATTGTCTAAC-3’ |
| pBRET-9-F | GCTGGTAATGCAACAGGCGGCGGAGGATCTATGAC-3’ |
| pBRET-9-R | TGTTGCATTACCAGCTTGGAAAGTAACACCTGAGCATTGTCTAAC-3’ |
| pBRET-10-F | GTAGCCACTGTACAGAGTGGTTTTAGAAAAATGGCATTCCCATCTGG-3’ |
| pBRET-10-R | CTGTACAGTGGCTACAGATCCTCCGCCGCCCTTG-3’ |

Table S2. The protein sequences of BRET-based self-cleaving biosensors.

| pBRET biosensors | Protein sequences^1^ |
| --- | --- |
| pBRET-1 | MDYKDDDDKMVSKGEELFTGVVPILVELDGDVNGHKFSVSGEGEGDATYGKLTLKFICTTGKLPVPWPTLVTTLSYGVQCFSRYPDHMKQHDFFKSAMPEGYVQERTIFFKDDGNYKTRAEVKFEGDTLVNRIELKGIDFKEDGNILGHKLEYNYNSHNVYIMADKQKNGIKVNFKIRHNIEDGSVQLADHYQQNTPIGDGPVLLPDNHYLSTQSALSKDPNEKRDHMVLLEFVTAAGITLGMDELYKGGGGSSAVLQSGFRKMAFPSGKVEGCMVQVTCGTTTLNGLWLDDVVYCPRHVICTSEDMLNPNYEDLLIRKSNHNFLVQAGNVQLRVIGHSMQNCVLKLKVDTANPKTPKYKFVRIQPGQTFSVLACYNGSPSGVYQCAMRPNFTIKGSFLNGSCGSVGFNIDYDCVSFCYMHHMELPTGVHAGTDLEGNFYGPFVDRQTAQAAGTDTTITVNVLAWLYAAVINGDRWFLNRFTTTLNDFNLVAMKYNYEPLTQDHVDILGPLSAQTGIAVLDMCASLKELLQNGMNGRTILGSALLEDEFTPFDVVRQCSGVTFQSAVKRGGGGSMTSKVYDPEQRKRMITGPQWWARCKQMNVLDSFINYYDSEKHAENAVIFLHGNATSSYLWRHVVPHIEPVARCIIPDLIGMGKSGKSGNGSYRLLDHYKYLTAWFELLNLPKKIIFVGHDWGAALAFHYAYEHQDRIKAIVHMESVVDVIESWDEWPDIEEDIALIKSEEGEKMVLENNFFVETVLPSKIMRKLEPEEFAAYLEPFKEKGEVRRPTLSWPREIPLVKGGKPDVVQIVRNYNAYLRASDDLPKLFIESDPGFFSNAIVEGAKKFPNTEFVKVKGLHFLQEDAPDEMGKYIKSFVERVLKNEQLEGSYPYDVPDYA* |
| pBRET-2 | MDYKDDDDKMVSKGEELFTGVVPILVELDGDVNGHKFSVSGEGEGDATYGKLTLKFICTTGKLPVPWPTLVTTLSYGVQCFSRYPDHMKQHDFFKSAMPEGYVQERTIFFKDDGNYKTRAEVKFEGDTLVNRIELKGIDFKEDGNILGHKLEYNYNSHNVYIMADKQKNGIKVNFKIRHNIEDGSVQLADHYQQNTPIGDGPVLLPDNHYLSTQSALSKDPNEKRDHMVLLEFVTAAGITLGMDELYKGGGGSGVTFQSGFRKMAFPSGKVEGCMVQVTCGTTTLNGLWLDDVVYCPRHVICTSEDMLNPNYEDLLIRKSNHNFLVQAGNVQLRVIGHSMQNCVLKLKVDTANPKTPKYKFVRIQPGQTFSVLACYNGSPSGVYQCAMRPNFTIKGSFLNGSCGSVGFNIDYDCVSFCYMHHMELPTGVHAGTDLEGNFYGPFVDRQTAQAAGTDTTITVNVLAWLYAAVINGDRWFLNRFTTTLNDFNLVAMKYNYEPLTQDHVDILGPLSAQTGIAVLDMCASLKELLQNGMNGRTILGSALLEDEFTPFDVVRQCSGVTFQSAVKRGGGGSMTSKVYDPEQRKRMITGPQWWARCKQMNVLDSFINYYDSEKHAENAVIFLHGNATSSYLWRHVVPHIEPVARCIIPDLIGMGKSGKSGNGSYRLLDHYKYLTAWFELLNLPKKIIFVGHDWGAALAFHYAYEHQDRIKAIVHMESVVDVIESWDEWPDIEEDIALIKSEEGEKMVLENNFFVETVLPSKIMRKLEPEEFAAYLEPFKEKGEVRRPTLSWPREIPLVKGGKPDVVQIVRNYNAYLRASDDLPKLFIESDPGFFSNAIVEGAKKFPNTEFVKVKGLHFLQEDAPDEMGKYIKSFVERVLKNEQLEGSYPYDVPDYA* |
| pBRET-3 | MDYKDDDDKMVSKGEELFTGVVPILVELDGDVNGHKFSVSGEGEGDATYGKLTLKFICTTGKLPVPWPTLVTTLSYGVQCFSRYPDHMKQHDFFKSAMPEGYVQERTIFFKDDGNYKTRAEVKFEGDTLVNRIELKGIDFKEDGNILGHKLEYNYNSHNVYIMADKQKNGIKVNFKIRHNIEDGSVQLADHYQQNTPIGDGPVLLPDNHYLSTQSALSKDPNEKRDHMVLLEFVTAAGITLGMDELYKGGGGSHTVLQSGFRKMAFPSGKVEGCMVQVTCGTTTLNGLWLDDVVYCPRHVICTSEDMLNPNYEDLLIRKSNHNFLVQAGNVQLRVIGHSMQNCVLKLKVDTANPKTPKYKFVRIQPGQTFSVLACYNGSPSGVYQCAMRPNFTIKGSFLNGSCGSVGFNIDYDCVSFCYMHHMELPTGVHAGTDLEGNFYGPFVDRQTAQAAGTDTTITVNVLAWLYAAVINGDRWFLNRFTTTLNDFNLVAMKYNYEPLTQDHVDILGPLSAQTGIAVLDMCASLKELLQNGMNGRTILGSALLEDEFTPFDVVRQCSGVTFQSAVKRGGGGSMTSKVYDPEQRKRMITGPQWWARCKQMNVLDSFINYYDSEKHAENAVIFLHGNATSSYLWRHVVPHIEPVARCIIPDLIGMGKSGKSGNGSYRLLDHYKYLTAWFELLNLPKKIIFVGHDWGAALAFHYAYEHQDRIKAIVHMESVVDVIESWDEWPDIEEDIALIKSEEGEKMVLENNFFVETVLPSKIMRKLEPEEFAAYLEPFKEKGEVRRPTLSWPREIPLVKGGKPDVVQIVRNYNAYLRASDDLPKLFIESDPGFFSNAIVEGAKKFPNTEFVKVKGLHFLQEDAPDEMGKYIKSFVERVLKNEQLEGSYPYDVPDYA* |
| pBRET-4 | MDYKDDDDKMVSKGEELFTGVVPILVELDGDVNGHKFSVSGEGEGDATYGKLTLKFICTTGKLPVPWPTLVTTLSYGVQCFSRYPDHMKQHDFFKSAMPEGYVQERTIFFKDDGNYKTRAEVKFEGDTLVNRIELKGIDFKEDGNILGHKLEYNYNSHNVYIMADKQKNGIKVNFKIRHNIEDGSVQLADHYQQNTPIGDGPVLLPDNHYLSTQSALSKDPNEKRDHMVLLEFVTAAGITLGMDELYKGGGGSFTRLQSGFRKMAFPSGKVEGCMVQVTCGTTTLNGLWLDDVVYCPRHVICTSEDMLNPNYEDLLIRKSNHNFLVQAGNVQLRVIGHSMQNCVLKLKVDTANPKTPKYKFVRIQPGQTFSVLACYNGSPSGVYQCAMRPNFTIKGSFLNGSCGSVGFNIDYDCVSFCYMHHMELPTGVHAGTDLEGNFYGPFVDRQTAQAAGTDTTITVNVLAWLYAAVINGDRWFLNRFTTTLNDFNLVAMKYNYEPLTQDHVDILGPLSAQTGIAVLDMCASLKELLQNGMNGRTILGSALLEDEFTPFDVVRQCSGVTFQSAVKRGGGGSMTSKVYDPEQRKRMITGPQWWARCKQMNVLDSFINYYDSEKHAENAVIFLHGNATSSYLWRHVVPHIEPVARCIIPDLIGMGKSGKSGNGSYRLLDHYKYLTAWFELLNLPKKIIFVGHDWGAALAFHYAYEHQDRIKAIVHMESVVDVIESWDEWPDIEEDIALIKSEEGEKMVLENNFFVETVLPSKIMRKLEPEEFAAYLEPFKEKGEVRRPTLSWPREIPLVKGGKPDVVQIVRNYNAYLRASDDLPKLFIESDPGFFSNAIVEGAKKFPNTEFVKVKGLHFLQEDAPDEMGKYIKSFVERVLKNEQLEGSYPYDVPDYA* |
| pBRET-5 | MDYKDDDDKMVSKGEELFTGVVPILVELDGDVNGHKFSVSGEGEGDATYGKLTLKFICTTGKLPVPWPTLVTTLSYGVQCFSRYPDHMKQHDFFKSAMPEGYVQERTIFFKDDGNYKTRAEVKFEGDTLVNRIELKGIDFKEDGNILGHKLEYNYNSHNVYIMADKQKNGIKVNFKIRHNIEDGSVQLADHYQQNTPIGDGPVLLPDNHYLSTQSALSKDPNEKRDHMVLLEFVTAAGITLGMDELYKGGGGSGVTFQSGFRKMAFPSGKVEGCMVQVTCGTTTLNGLWLDDVVYCPRHVICTSEDMLNPNYEDLLIRKSNHNFLVQAGNVQLRVIGHSMQNCVLKLKVDTANPKTPKYKFVRIQPGQTFSVLACYNGSPSGVYQCAMRPNFTIKGSFLNGSCGSVGFNIDYDCVSFCYMHHMELPTGVHAGTDLEGNFYGPFVDRQTAQAAGTDTTITVNVLAWLYAAVINGDRWFLNRFTTTLNDFNLVAMKYNYEPLTQDHVDILGPLSAQTGIAVLDMCASLKELLQNGMNGRTILGSALLEDEFTPFDVVRQCSGVTFQSLENVGGGGSMTSKVYDPEQRKRMITGPQWWARCKQMNVLDSFINYYDSEKHAENAVIFLHGNATSSYLWRHVVPHIEPVARCIIPDLIGMGKSGKSGNGSYRLLDHYKYLTAWFELLNLPKKIIFVGHDWGAALAFHYAYEHQDRIKAIVHMESVVDVIESWDEWPDIEEDIALIKSEEGEKMVLENNFFVETVLPSKIMRKLEPEEFAAYLEPFKEKGEVRRPTLSWPREIPLVKGGKPDVVQIVRNYNAYLRASDDLPKLFIESDPGFFSNAIVEGAKKFPNTEFVKVKGLHFLQEDAPDEMGKYIKSFVERVLKNEQLEGSYPYDVPDYA* |
| pBRET-6 | MDYKDDDDKMVSKGEELFTGVVPILVELDGDVNGHKFSVSGEGEGDATYGKLTLKFICTTGKLPVPWPTLVTTLSYGVQCFSRYPDHMKQHDFFKSAMPEGYVQERTIFFKDDGNYKTRAEVKFEGDTLVNRIELKGIDFKEDGNILGHKLEYNYNSHNVYIMADKQKNGIKVNFKIRHNIEDGSVQLADHYQQNTPIGDGPVLLPDNHYLSTQSALSKDPNEKRDHMVLLEFVTAAGITLGMDELYKGGGGSGVTFQSGFRKMAFPSGKVEGCMVQVTCGTTTLNGLWLDDVVYCPRHVICTSEDMLNPNYEDLLIRKSNHNFLVQAGNVQLRVIGHSMQNCVLKLKVDTANPKTPKYKFVRIQPGQTFSVLACYNGSPSGVYQCAMRPNFTIKGSFLNGSCGSVGFNIDYDCVSFCYMHHMELPTGVHAGTDLEGNFYGPFVDRQTAQAAGTDTTITVNVLAWLYAAVINGDRWFLNRFTTTLNDFNLVAMKYNYEPLTQDHVDILGPLSAQTGIAVLDMCASLKELLQNGMNGRTILGSALLEDEFTPFDVVRQCSGVTFQAIASEGGGGSMTSKVYDPEQRKRMITGPQWWARCKQMNVLDSFINYYDSEKHAENAVIFLHGNATSSYLWRHVVPHIEPVARCIIPDLIGMGKSGKSGNGSYRLLDHYKYLTAWFELLNLPKKIIFVGHDWGAALAFHYAYEHQDRIKAIVHMESVVDVIESWDEWPDIEEDIALIKSEEGEKMVLENNFFVETVLPSKIMRKLEPEEFAAYLEPFKEKGEVRRPTLSWPREIPLVKGGKPDVVQIVRNYNAYLRASDDLPKLFIESDPGFFSNAIVEGAKKFPNTEFVKVKGLHFLQEDAPDEMGKYIKSFVERVLKNEQLEGSYPYDVPDYA* |
| pBRET-7 | MDYKDDDDKMVSKGEELFTGVVPILVELDGDVNGHKFSVSGEGEGDATYGKLTLKFICTTGKLPVPWPTLVTTLSYGVQCFSRYPDHMKQHDFFKSAMPEGYVQERTIFFKDDGNYKTRAEVKFEGDTLVNRIELKGIDFKEDGNILGHKLEYNYNSHNVYIMADKQKNGIKVNFKIRHNIEDGSVQLADHYQQNTPIGDGPVLLPDNHYLSTQSALSKDPNEKRDHMVLLEFVTAAGITLGMDELYKGGGGSGVTFQSGFRKMAFPSGKVEGCMVQVTCGTTTLNGLWLDDVVYCPRHVICTSEDMLNPNYEDLLIRKSNHNFLVQAGNVQLRVIGHSMQNCVLKLKVDTANPKTPKYKFVRIQPGQTFSVLACYNGSPSGVYQCAMRPNFTIKGSFLNGSCGSVGFNIDYDCVSFCYMHHMELPTGVHAGTDLEGNFYGPFVDRQTAQAAGTDTTITVNVLAWLYAAVINGDRWFLNRFTTTLNDFNLVAMKYNYEPLTQDHVDILGPLSAQTGIAVLDMCASLKELLQNGMNGRTILGSALLEDEFTPFDVVRQCSGVTFQSKMSDGGGGSMTSKVYDPEQRKRMITGPQWWARCKQMNVLDSFINYYDSEKHAENAVIFLHGNATSSYLWRHVVPHIEPVARCIIPDLIGMGKSGKSGNGSYRLLDHYKYLTAWFELLNLPKKIIFVGHDWGAALAFHYAYEHQDRIKAIVHMESVVDVIESWDEWPDIEEDIALIKSEEGEKMVLENNFFVETVLPSKIMRKLEPEEFAAYLEPFKEKGEVRRPTLSWPREIPLVKGGKPDVVQIVRNYNAYLRASDDLPKLFIESDPGFFSNAIVEGAKKFPNTEFVKVKGLHFLQEDAPDEMGKYIKSFVERVLKNEQLEGSYPYDVPDYA* |
| pBRET-8 | MDYKDDDDKMVSKGEELFTGVVPILVELDGDVNGHKFSVSGEGEGDATYGKLTLKFICTTGKLPVPWPTLVTTLSYGVQCFSRYPDHMKQHDFFKSAMPEGYVQERTIFFKDDGNYKTRAEVKFEGDTLVNRIELKGIDFKEDGNILGHKLEYNYNSHNVYIMADKQKNGIKVNFKIRHNIEDGSVQLADHYQQNTPIGDGPVLLPDNHYLSTQSALSKDPNEKRDHMVLLEFVTAAGITLGMDELYKGGGGSGVTFQSGFRKMAFPSGKVEGCMVQVTCGTTTLNGLWLDDVVYCPRHVICTSEDMLNPNYEDLLIRKSNHNFLVQAGNVQLRVIGHSMQNCVLKLKVDTANPKTPKYKFVRIQPGQTFSVLACYNGSPSGVYQCAMRPNFTIKGSFLNGSCGSVGFNIDYDCVSFCYMHHMELPTGVHAGTDLEGNFYGPFVDRQTAQAAGTDTTITVNVLAWLYAAVINGDRWFLNRFTTTLNDFNLVAMKYNYEPLTQDHVDILGPLSAQTGIAVLDMCASLKELLQNGMNGRTILGSALLEDEFTPFDVVRQCSGVTFQNNELSGGGGSMTSKVYDPEQRKRMITGPQWWARCKQMNVLDSFINYYDSEKHAENAVIFLHGNATSSYLWRHVVPHIEPVARCIIPDLIGMGKSGKSGNGSYRLLDHYKYLTAWFELLNLPKKIIFVGHDWGAALAFHYAYEHQDRIKAIVHMESVVDVIESWDEWPDIEEDIALIKSEEGEKMVLENNFFVETVLPSKIMRKLEPEEFAAYLEPFKEKGEVRRPTLSWPREIPLVKGGKPDVVQIVRNYNAYLRASDDLPKLFIESDPGFFSNAIVEGAKKFPNTEFVKVKGLHFLQEDAPDEMGKYIKSFVERVLKNEQLEGSYPYDVPDYA* |
| pBRET-9 | MDYKDDDDKMVSKGEELFTGVVPILVELDGDVNGHKFSVSGEGEGDATYGKLTLKFICTTGKLPVPWPTLVTTLSYGVQCFSRYPDHMKQHDFFKSAMPEGYVQERTIFFKDDGNYKTRAEVKFEGDTLVNRIELKGIDFKEDGNILGHKLEYNYNSHNVYIMADKQKNGIKVNFKIRHNIEDGSVQLADHYQQNTPIGDGPVLLPDNHYLSTQSALSKDPNEKRDHMVLLEFVTAAGITLGMDELYKGGGGSGVTFQSGFRKMAFPSGKVEGCMVQVTCGTTTLNGLWLDDVVYCPRHVICTSEDMLNPNYEDLLIRKSNHNFLVQAGNVQLRVIGHSMQNCVLKLKVDTANPKTPKYKFVRIQPGQTFSVLACYNGSPSGVYQCAMRPNFTIKGSFLNGSCGSVGFNIDYDCVSFCYMHHMELPTGVHAGTDLEGNFYGPFVDRQTAQAAGTDTTITVNVLAWLYAAVINGDRWFLNRFTTTLNDFNLVAMKYNYEPLTQDHVDILGPLSAQTGIAVLDMCASLKELLQNGMNGRTILGSALLEDEFTPFDVVRQCSGVTFQAGNATGGGGSMTSKVYDPEQRKRMITGPQWWARCKQMNVLDSFINYYDSEKHAENAVIFLHGNATSSYLWRHVVPHIEPVARCIIPDLIGMGKSGKSGNGSYRLLDHYKYLTAWFELLNLPKKIIFVGHDWGAALAFHYAYEHQDRIKAIVHMESVVDVIESWDEWPDIEEDIALIKSEEGEKMVLENNFFVETVLPSKIMRKLEPEEFAAYLEPFKEKGEVRRPTLSWPREIPLVKGGKPDVVQIVRNYNAYLRASDDLPKLFIESDPGFFSNAIVEGAKKFPNTEFVKVKGLHFLQEDAPDEMGKYIKSFVERVLKNEQLEGSYPYDVPDYA* |
| pBRET-10 | MDYKDDDDKMVSKGEELFTGVVPILVELDGDVNGHKFSVSGEGEGDATYGKLTLKFICTTGKLPVPWPTLVTTLSYGVQCFSRYPDHMKQHDFFKSAMPEGYVQERTIFFKDDGNYKTRAEVKFEGDTLVNRIELKGIDFKEDGNILGHKLEYNYNSHNVYIMADKQKNGIKVNFKIRHNIEDGSVQLADHYQQNTPIGDGPVLLPDNHYLSTQSALSKDPNEKRDHMVLLEFVTAAGITLGMDELYKGGGGSVATVQSGFRKMAFPSGKVEGCMVQVTCGTTTLNGLWLDDVVYCPRHVICTSEDMLNPNYEDLLIRKSNHNFLVQAGNVQLRVIGHSMQNCVLKLKVDTANPKTPKYKFVRIQPGQTFSVLACYNGSPSGVYQCAMRPNFTIKGSFLNGSCGSVGFNIDYDCVSFCYMHHMELPTGVHAGTDLEGNFYGPFVDRQTAQAAGTDTTITVNVLAWLYAAVINGDRWFLNRFTTTLNDFNLVAMKYNYEPLTQDHVDILGPLSAQTGIAVLDMCASLKELLQNGMNGRTILGSALLEDEFTPFDVVRQCSGVTFQNNELSGGGGSMTSKVYDPEQRKRMITGPQWWARCKQMNVLDSFINYYDSEKHAENAVIFLHGNATSSYLWRHVVPHIEPVARCIIPDLIGMGKSGKSGNGSYRLLDHYKYLTAWFELLNLPKKIIFVGHDWGAALAFHYAYEHQDRIKAIVHMESVVDVIESWDEWPDIEEDIALIKSEEGEKMVLENNFFVETVLPSKIMRKLEPEEFAAYLEPFKEKGEVRRPTLSWPREIPLVKGGKPDVVQIVRNYNAYLRASDDLPKLFIESDPGFFSNAIVEGAKKFPNTEFVKVKGLHFLQEDAPDEMGKYIKSFVERVLKNEQLEGSYPYDVPDYA* |

^1^Protein sequences of different components in the biosensors were identified as follows: green, GFP2; orange, SARS-CoV-2 3CLpro; blue, RLuc8; underscored, 3CLpro cleavage sequences. DYKDDDDK and YPYDVPDYA are the sequences of FLAG tag and HA tag, respectively.
